# Disruption of Immune Responses By Type I Diabetes Exacerbates SARS-CoV-2 Mediated Lung Injury

**DOI:** 10.1101/2024.05.31.596857

**Authors:** Sara Kass-Gergi, Gan Zhao, Joanna Wong, Aaron I. Weiner, Stephanie Adams Tzivelekidis, Maria E. Gentile, Meryl Mendoza, Nicolas P. Holcomb, Xinyuan Li, Madeline Singh, Andrew E. Vaughan

**Author notes:** Corresponding authors, to whom correspondence should be addressed: Andrew E. Vaughan. The authors have declared that no conflict of interest exists.

## Abstract

COVID-19 commonly presents as pneumonia, with those most severely affected progressing to respiratory failure. Patient responses to SARS-CoV-2 infection are varied, with comorbidities acting as major contributors to varied outcomes. Focusing on one such major comorbidity, we assessed whether pharmacological induction of Type I Diabetes Mellitus (T1DM) would increase the severity of lung injury in a murine model of COVID-19 pneumonia utilizing wild type mice infected with mouse-adapted SARS-CoV-2. Hyperglycemic mice exhibited increased weight loss and reduced blood oxygen saturation in comparison to their euglycemic counterparts, suggesting that these animals indeed experienced more severe lung injury. Transcriptomic analysis revealed a significant impairment of the adaptive immune response in the lungs of diabetic mice compared to those of control. In order to expand the limited options available for tissue analysis due to biosafety restrictions, we also employed a novel technique to digest highly fixed tissue into a single cell suspension, which allowed for flow cytometric analysis as well as single cell RNA sequencing. Flow immunophenotyping and scRNA-Seq confirmed impaired recruitment of T cells into the lungs of T1DM animals. Additionally, scRNA-Seq revealed a distinct, highly inflammatory macrophage profile in the diabetic cohort that correlates with the more severe infection these mice experienced clinically, allowing insight into a possible mechanism for this phenomenon. Recognizing the near certainty that respiratory viruses will continue to present significant public health concerns for the foreseeable future, our study provides key insights into how T1DM results in a much more severe infection and identifies possible targets to ameliorate comorbidity-associated severe disease.

**NEW AND NOTEWORTHY:** We define the exacerbating effects of Type I Diabetes Mellitus (T1DM) on COVID-19 pneumonia severity in mice. Hyperglycemic mice experienced increased weight loss and reduced oxygen saturation. Transcriptomic analysis revealed impaired immune responses in diabetic mice, while flow cytometry and single-cell RNA sequencing confirmed reduced T cell recruitment and an inflammatory macrophage profile. Additionally, we introduced a novel technique for tissue analysis, enabling flow cytometric analysis and single-cell RNA sequencing on highly fixed tissue samples.

## INTRODUCTION

SARS-CoV-2 infection / COVID-19 commonly presents as pneumonia, with those most severely affected progressing to Acute Respiratory Distress Syndrome (ARDS), resulting in a mortality rate as high at 40%(1). Patient responses to SARS-CoV-2 infection are heterogeneous, ranging from asymptomatic to ARDS to initially mild disease later manifesting chronic sequelae (“long COVID”)(1, 2). At least some of this heterogeneity is due to patient risk factors and comorbidities, with metabolic diseases including type I and type 2 diabetes mellitus, as well as obesity and hyperlipidemia, significantly increasing the odds of severe disease(2–5). Despite numerous theories, the molecular basis by which these comorbidities promote disease progression remains unclear.

Type I diabetes (T1DM) is an autoimmune disease caused by the destruction of the beta cells of the pancreas responsible for making insulin(6). While there has been a plethora of research into the underlying autoimmunity responsible for the onset of T1DM, the subsequent impact of hyperglycemia and the systemic diabetic phenotype on immune function remain understudied(7). Though studies in both type I and type 2 DM have shown that hyperglycemia itself has a negative impact on the immune system, it does not seem to be the only factor mediating this dysfunction(7, 8). In addition to COVID, patients with diabetes have been found to have worse outcomes with other respiratory viruses, such as influenza and respiratory syncytial virus(9, 10).

In this study, we sought to explore the effect of T1DM and hyperglycemia on the severity and course of SARS-CoV-2 infection. We utilize a well-established murine model of T1DM(11) and a mouse adapted strain of SARS-CoV-2 known to induce significant pulmonary disease(12), and we observed that hyperglycemia does in fact confer a much more severe and morbid SARS-CoV-2 infection. Furthermore, through transcriptomic and cytometric analyses, we identified a critical deficiency in the adaptive immune system that likely explains at least some of the more deleterious effects of infection on the lungs of T1DM mice including impaired viral clearance. Employing single cell transcriptomics, we demonstrate that upon infection, diabetic animals with impaired adaptive immunity exhibit an enhanced proinflammatory phenotype of alveolar macrophages that likely serve as a compensatory, if dysfunctional, response leading to increased morbidity. In addition, as a result of innovations necessary for analysis of tissue from Animal Biosafety Level 3 (ABSL-3) conditions, we present here a novel protocol that allows for thorough and quantitative immunophenotypic analysis of fully fixed tissues, even those which had been previously embedded for cryosectioning. This technique will allow for greatly expanded analyses of fixed tissues broadly from high biosafety level experiments and archived samples.

## METHODS

### Mice

Wild-type C57BL/6 male mice (Jackson) were housed in housed in the Animal Biosafety Level 1 Facility at the University of Pennsylvania under the standard dark/light cycle, ambient temperature, humidity, and within the Allentown individual ventilated BCU2 caging system. Animals were 8 weeks of age and littermates, randomly assigned to each group. All animal experiments were carried out under the guidelines set by the University of Pennsylvania’s Institutional Animal Care and Use Committees and followed all National Institutes of Health (NIH) Office of Laboratory Animal Welfare regulations.

### Streptozotocin-induced Type 1 diabetes

Mice were housed in Animal Biosafety Level 2 facility. Adult mice were randomly assigned to receive 150mg/kg STZ in sodium citrate buffer or sodium citrate buffer as vehicle control via intraperitoneal injection(11) on day -28 and were allowed to recover for four weeks before further experimentation. Sucrose was added to their water (30% sucrose by volume) for 5 days post injection to support possible hypoglycemia as a result of the pancreatic beta cells releasing stored insulin subsequent to their destruction. Blood glucose levels were checked on days -18 and -7 after 6 hours of fasting using tail vein blood samples and Contour Next EZ Blood Glucose Monitoring System. Mice who had received STZ but did not meet criteria for hyperglycemia (fasting blood glucose >250mg/dL) were excluded.

### Mouse-adapted SARS-CoV-2 injury

Mice were housed in the Animal Biosafety Level 3 Facility at the University of Pennsylvania under the standard dark/light cycle, ambient temperature, humidity, and within the Allentown individual ventilated BCU2 caging system. All work was in adherence to the University of Pennsylvania’s approved ABSL-3 IBC and IACUC protocols. For some experiments (whole lung bulk RNA sequencing, Figure 2) mice were randomly assigned to one of two treatment groups: mock infected and infected. Mice were anesthetized with ketamine-xylazine and infected with 10^4^ PFU’s of mouse-adapted SARS-CoV-2 (SARS2-N501Y_MA30_) intranasally in 40 µl DMEM. The virus was propagated from the MA30 stock virus (a generous gift from Stanley Perleman) in TMPRSS2-expressing Vero cells(12). The TMPRSS2 Vero cells were grown in Dulbecco’s modified Eagle’s medium DMEM, supplemented with 10% FBS. The virus was sequenced after propagation and found to match the input strain. Mice were monitored daily and scored for clinical disease and body weight was measured. At clinical or experimental endpoint (7 days post-infection), mice were anesthetized by isoflurane and necropsied. All tissues were inactivated and removed from the Animal Biosafety Level 3 facility in accordance with University of Pennsylvania IBC protocol.

### Whole lung bulk RNA sequencing

Lung tissue was placed in TRIzol® reagent (ThermoFisher, Catalog no. 15596018) RNA was extracted using Direct-zol RNA Miniprep Plus kit from Zymo. cDNA synthesis, sequencing library generation, and sequencing was performed on an Illumina HiSeq platform by GENEWIZ Co. Ltd. Raw data (raw reads) in fastq.gz format were processed through a general pipeline as describe previously (13). Reads were aligned to the mm10 mouse genome using Kallisto and imported into R Studio for analysis via the TxImport package. Data was then normalized using the trimmed mean of M values normalization method in the EdgeR package. Mean-variance trend fitting, linear modeling, and Bayesian statistics for differential gene expression analysis were performed using the Voom, LmFit, and eBayes functions, respectively, of the Limma package, yielding differentially expressed genes between uninfected, infected, euglycemic, and diabetic groups. Volcano plots were created using the OmicStudio tools at https://www.omicstudio.cn/tool. All detectable genes derived from RNA-seq were used for gene set enrichment analysis (GSEA) using the Molecular Signatures Database (MSigDB) C2: curated gene sets according to the standard GSEA user guide (http://www.broadinstitute.org/gsea/doc/GSEAUserGuideFrame.html).

### RNA isolation and qPCR

Total RNA was extracted by Direct-zol RNA Miniprep Plus kit according to the manufacturer’s instruction (Zymo, Catalog no. R2072) and then reverse-transcribed into complementary DNA using the iScript Reverse Transcription Supermix (Bio-Rad, Catalog no. 1708841). qPCR was performed using a PowerUp SYBR Green Master Mix and standard protocols on an Applied Biosystems QuantStudio 6 Real-Time PCR System (Thermo Fisher Scientific). RPL19 was used to normalize RNA isolated from mouse samples. The 2^−ΔΔ^Ct comparative method was used to analyze expression levels. The primers used are listed in Supplementary Table 1.

### Fixed tissue single cell digest

Given the requirements for ABSL-3 safety precautions, the tissue harvested from these mice needs to be fixed in paraformaldehyde for at least 24 hours prior to further handling. Logistically, the digestion of tissue into a single cell suspension prior to the required fixation step was impossible due to time and staff constraints. To facilitate these experiments, we adapted a novel technique originally developed for the analysis of fixed tissue for scRNA-seq, Tissue Fixation & Dissociation for Chromium Fixed RNA Profiling (10x Genomics, Protocol CG000553). Tissue was harvested in the ABSL-3 facility by trained personnel and minced into square millimeter sized pieces before being fixed for 24 hours at 4°C. Samples dedicated to scRNA sequencing were fixed with Fix & Perm Buffer (10x Genomics, PN-2000517) as per manufacturer’s protocol while samples for flow cytometry were fixed in 4% PFA. Samples were then washed with PBS and the fixation reaction was quenched. scRNA samples were quenched with Quench Buffer (10x Genomics, PN-2000516) as per manufacturer’s protocol, while flow cytometry samples were quenched in solution of 50mM glycine dissolved in PBS. Tissue was then dissociated into a single cell suspension shaking vigorously at 37°C for 1 hour in Dissociation Solution made with Liberase^TM^ TL (concentration 0.2mg/mL, Millipore Sigma, Catalog no. 5401020001) as per manufacturer’s protocol. Tissue was further triturated with a P1000 pipette tip 15-20 times before being filtered through a 40µM filter, which was then washed with PBS. The samples were quenched again with Quench Buffer and cells were then counted by hemocytometer before proceeding to scRNA sequencing or flow cytometry. Samples were stored at 4°C for up to 1 week before use. Samples designated for scRNA sequencing were incubated in Quenching Buffer and Enhancer (10x Genomics, PN-2000482) as per manufacturer’s protocol.

### Histology

Lung tissues were fixed in 4% PFA for 24 hours at room temperature in accordance with the ABSL-3 safety protocols, rinsed three times with PBS at room temperature, incubated in 30% sucrose overnight at 4°C, and transferred to 15% sucrose 50% optimal cutting temperature (OCT) compound at room temperature for 4 hours. Fixed tissues were transferred to an embedding mold filled with OCT compound, frozen on dry ice, and stored at -80°C. Frozen sections (7 µm thick) were cut at -20°C, and slides were kept at -20°C until use. Sections were stained with Hematoxylin and Eosin Stain Kit (Vector Laboratories, Catalog no. H-3502) according to the manufacture’s protocol and then imaged with a Leica DMi8 microscope.

### Fluorescence activated cell analysis

Whole lung single-cell suspensions from mice on day 7 post-MA30-SARS-CoV2 infected lungs were prepared as above. Single-cell suspensions were blocked in FACS buffer containing 1:50 mouse TruStain FcX for 10 minutes at room temperature. The cell suspension was stained using Brilliant Violet 785™ anti-mouse CD45 antibody (1:100, Biolegend, Clone 30-F11), Brilliant Violet 421™ anti-mouse F4/80 antibody (1:100; Biolegend, Clone BM8), PE/Cyanine5 anti-mouse CD3ε antibody (1:200, Biolegend, Clone 145-2C11), Brilliant Violet 785™ anti-mouse CD4 antibody (1:50, Biolegend, Clone GK1.5), APC anti-mouse CD8a Recombinant antibody (1:100, Biolegend, Clone QA17A07), PE anti-mouse Ly-6G antibody (1:200; Biolegend, Clone 1A8), PE/Cyanine7 anti-mouse CD64 (FcγRI) antibody (1:200, Biolegend, Clone X54-5/7.1), Brilliant Violet 711™ anti-mouse/human CD11b antibody (1:200; Biolegend, Clone M1/70), and FITC anti-mouse/human CD45R/B220 antibody (1:200, Biolegend, Clone RA3-6B2) overnight at 4°C. Stained cells and “fluorescence minus one” (FMO) controls were then resuspended in FACS buffer. All flow analyses were performed on BD LSRFortessa (BD Biosciences).

### Single Cell RNA Sequencing Library Construction

Whole lung single-cell suspensions from mice on day 7 post-MA30-SARS-CoV2 infected lungs was prepared as above and were used for sequencing. Single-cell sequencing libraries were created according to 10x Genomics Chromium Fixed RNA Profiling Reagents (10x Genomics, PN-1000414) and associated user protocol (User Guide CG000477 Rev D). After fixation, the samples were first hybridized using mouse specific whole transcriptome probe pairs. Cell counts were verified using a LogosBio Luna-FL automated cell counter(1.05e6 cells•mL-1 and 2.8e6 cells•mL-1 for sample 708L and 719L, respectively). The cells were pelletized and re-suspended in a hybridization mixture prepared using 10x Genomics Hyb Buffer B (PN-2000483) and Enhancer (PN-2000482) according to the protocol. Mouse specific WTA Probes BC001 (10x Genomics, PN-2000703) were then added to each sample at room temperature. The samples were incubated at 42°C in a Bio-Rad C1000 Touch Thermal Cycler for 16 hours.

Following probe hybridization, each sample was washed in Post-Hyb Wash Buffer prepared according to the protocol using concentrated Post-Hyb Buffer (PN-2000533) and Enhancer (PN-2000482). The samples were then incubated 42°C, centrifuged for 5 minutes at 850 rcf and resuspended in Post-Hyb Buffer. This wash procedure was repeated for a total of three washes. Each sample was then filtered through a 30 µm Sysmex CellTrics filter into a clean 1.5mL Lo-Bind Eppendorf Tube. The cell concentration was then verified again using the LogosBio Luna-FL automated cell counter according to the manufacturers protocol (3.86e6 cells•mL-1 and 1.59e6 cells•mL-1 for sample 708L and 719L, respectively). Gel Beads-in-Emulsion (GEMs) were then generated using the 10x Next GEM Chip Q according to the manufacturer’s protocol and loaded onto a 10x Genomics Chromium X Chip Reader. Cells were loaded onto the chip targeting a total of 10,000 cells per sample. Upon completion of the chip being processed, the GEMs were transferred to new PCR strip tubes and incubated in a Bio-Rad C1000 Thermal Cycler according to the protocol. The samples were left in a 4°C refrigeration overnight.

The samples were then mixed with 10X Genomics proprietary recovery agent (PN-220016). The recovery agent is then removed and discarded after 2 minutes from resulting biphasic mixture. The remaining aqueous phase is then amplified according to the manufacturer’s protocol. A DNA cleanup is then performed on the amplified DNA using Beckman Coulter SPRIselect magnetic beads. Gene Expression libraries were then created according to the protocol, using 12 cycles of the appropriate PCR program supplied by the manufacturer. The samples then underwent another cleanup using SPRIselect magnetic beads. Libraries were quantified using the Invitrogen High Sensitivity DNA Qubit Assay and library quality was checked using the Agilent Tapestation 4200 controller and High Sensitivity D1000 Assay. The libraries were sequenced on an Illumina NextSeq 2000 using a P2 100 Cycle kit. A 2.05 nM pool was prepared, and the sequencing cartridge was loaded at 650 pM with 1µL of 1nM PhiX. A minimum of 10,000 read pairs were targeted for each sample with paired-end dual indexing. The parameters for each read were 28 cycles for Read 1, 10 cycles for the i7 index, 10 cycles for the i5 index, and 90 cycles for Read 2, per the protocol.

### Single cell transcriptomics

After sequencing, initial data processing was performed using Cellranger (v7.2.0). Cellranger mkfastq was used to generate demultiplexed FASTQ files from the raw sequencing data. Next, Cellranger count was used to align sequencing reads to the mouse reference genome (GRCm38) and generate single-cell gene barcode matrices. Post-processing and secondary analysis were performed using the Seurat package (v.4.0)(14). First, variable features across single cells in the dataset were identified by mean expression and dispersion. Identified variable features was then be used to perform a PCA. The dimensionally reduced data was used to cluster cells and visualize using a UMAP plot(15). Cell populations were identified by comparison to known marker genes for each cell type. Differential expression analysis between samples and populations within samples was performed using default variable with the command FindMarkers.

### Statistics

All statistical calculations were performed using GraphPad Prism 9. Unpaired two-tailed Student’s t-tests were used to ascertain statistical significance between two groups. For details on statistical analyses, tests used, size of n, definition of significance, and summaries of statistical outputs, see corresponding figure legend and the Results section.

### Data Availability

Single-cell RNA-sequencing data generated in this study will be made available via the NIH Gene Expression Omnibus (GEO) upon release of peer reviewed publication.

## RESULTS

### Hyperglycemia renders mice more susceptible to severe SARS-CoV2 infection and impairs recovery

The utilization of high-dose streptozotocin (STZ) is an effective and widely used technique for inducing T1DM in mice(11). This alkylating agent operates by selectively binding to and inducing toxicity of the pancreatic β-cells, which are responsible for insulin production. Consequently, this model closely replicates the pathophysiology of T1DM. We treated wild-type C57BL/6 mice with a single, high dose of STZ (Figure 1A) which resulted in the intended outcome of hyperglycemia after 10 days (Figure 1B). We then subjected hyperglycemic and vehicle-treated, euglycemic control mice to infection with MA30-SARS-CoV-2, a mouse-adapted strain recognized for inducing significant pulmonary disease. Specifically, T1DM-afflicted mice exhibited more pronounced weight loss, exacerbated oxygenation deficits, and a slower recovery trajectory in comparison to their euglycemic counterparts (Figure 1C, D). While not dramatic in either group, tissue damage was apparent in the lungs after infection as judged by hematoxylin and eosin staining (Figure 1F). Although these outcomes were apparent in both sexes, the differences were more stark in males (Supplementary Figure 1), so we utilized males for subsequent experiments unless otherwise noted. We performed qPCR analysis of RNA extracted from the whole lungs of both DM and non-DM mice at day 7 post infection to assess the level of SARS-CoV-2 nucleocapsid protein transcript as a substitute for viral plaque assays given the restrictions of ABSL-3 safety precautions. Critically, we noted that T1DM mice were markedly less effective in clearing the virus by the seventh day post-infection, while euglycemic mice had undetectable viral loads by this time point (Figure 1E). Intriguingly, and in contrast to recently published work(16), we did not observe any overt phenotype upon H1N1 influenza infection in these mice (Supplementary Figure 2). These findings not only emphasize the impact of diabetes on the severity of SARS-CoV-2 infection but also underscore the intricate interplay between metabolic disorders and viral pathogenesis and indicate that at least some components of increased morbidity of T1DM subjects result from an impaired ability to effectively clear SARS-CoV-2 virus.

**Figure 1.**
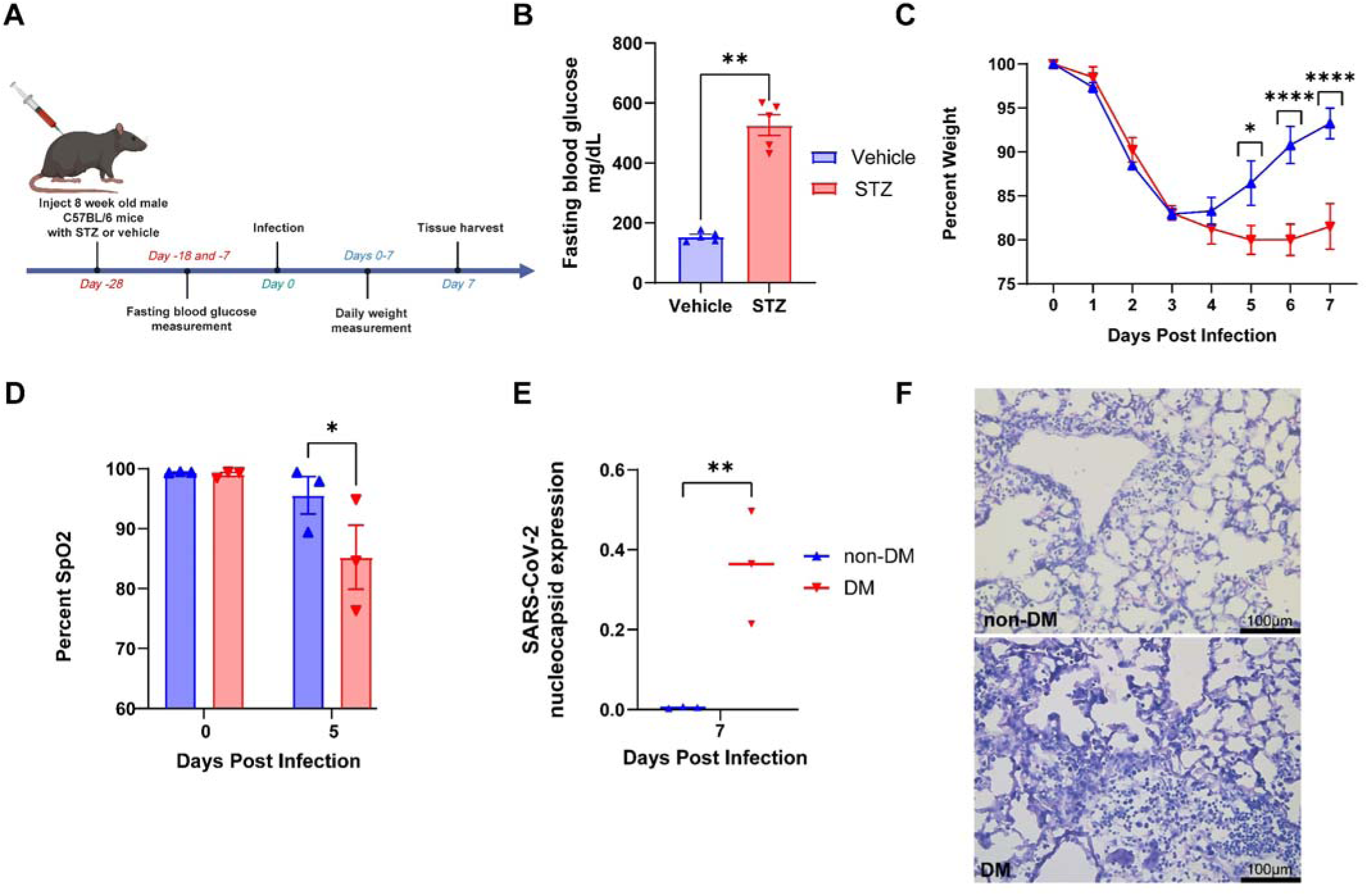
Hyperglycemia and T1DM renders mice more susceptible to severe SARS-CoV-2 infection. A. Schematic of experimental design for inducing T1DM in wild type mice. Figure made with Biorender. B. STZ effectively creates fasting hyperglycemia in wild type mice C. Time course of changes in body weight in male WT (non-DM) and T1DM mice after MA30SARS-CoV-2 infection, n = 4 to 7 per group. D. Comparison of capillary oxygen saturation of male non-DM and DM mice at day 0 (uninjured) and day 5 post infection, n = 3 per group. E. qPCR analysis of SARS-CoV-2 nucleocapsid protein expression in lungs of DM and non-DM mice harvested 7 days post infection, n = 3 per group. F. Representative H&E-stained sections of lung tissue harvested from non-DM and DM mice 7 days post infection with MA30SARS-CoV-2, scale bars 100 µm. Data in (B), (C), (D), and (E) are presented as Mean ± SEM, calculated using one-way analysis of variance (ANOVA), followed by Mann Whitney test (B), Sidak’s multiple comparison test (C), Uncorrected Fisher’s LSD (D), and Turkey’s multiple comparison test (E). *p < 0.05, **p < 0.01, and ****p < 0.0001.

**Figure 2.**
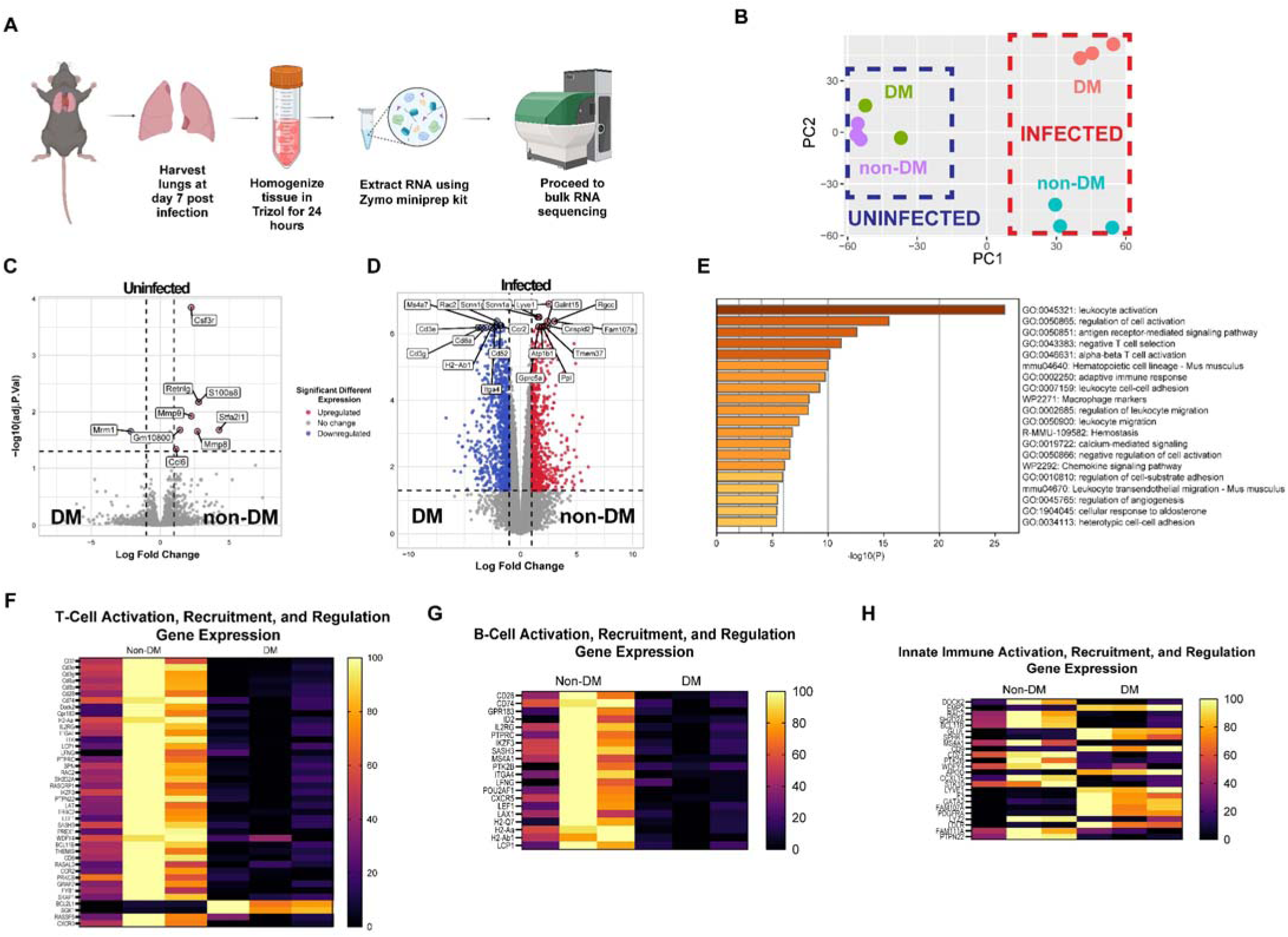
Whole lung RNA sequencing reveals an impaired adaptive immune response in DM mice infected with MA30-SARS-CoV-2. A. Schematic for experimental design of whole lung RNA sequencing. Figure made with Biorender. B. PCA plot of whole lung RNA sequencing of tissue harvested from non-DM, uninfected mice, DM, uninfected mice, non-DM, infected mice, and DM, infected mice, n =3 per group. C-D. Volcano plots of significantly differentially expressed genes (C) in the lungs of uninfected non-DM and DM mice (differential gene expression meeting significance = 9 genes) and (D) in the lungs of infected non-DM and DM mice (differential gene expression meeting significance = 1745 genes) A. The GO terms of differentially expressed genes in non-DM and DM mice lungs infected with MA30-SARS-CoV-2 were carried out by Metascape (36). F-H. Heatmap showing the top differentially expressed genes between non-DM and DM mice in relation to the activation, recruitment and regulation of (F) T-cells, (G) B-cells, and (H) innate immune cells.

### Whole lung RNA sequencing reveals deficiency in adaptive immune response in diabetic mice as a potential etiology for prolonged morbidity and impaired recovery

We sought to take an unbiased approach to hypothesis generation for underlying mechanisms to explain the differences in clinical course and viral clearance observed in T1DM animals. We thus employed whole lung, “bulk” RNA sequencing to further examine the transcriptomic changes. Importantly, while there were only minimal changes (9 genes) present between the two groups of uninfected mice, upon infection the number of differentially expressed genes increased sharply to >1500 (Figure 2C, D), reinforcing the dramatic changes in diabetic host response upon MA-SARS-CoV-2 infection. Further examination of the data highlighted a significant downregulation of genes specifically associated with the adaptive immune system in the lungs of DM mice, as illustrated in Figure 2E. This downregulation extended to genes crucial for the activation, recruitment, and proliferation of both T-cells (Figure 2F) and B-cells (Figure 2G). Interestingly, many genes associated with innate immunity were also differentially expressed, though with no clear pattern weighted toward either group of infected mice. Taken together, these results indicate that hyperglycemia / T1DM renders mice more susceptible to severe disease with MA30-SARS-CoV-2, and that impaired adaptive immunity appears to explain many components of the observed phenotype including the persistence of viral particles and relative downregulation of adaptive immune gene markers in T1DM animals upon infection.

### Development of a novel flow cytometric immunophenotyping strategy for fixed tissue corroborates impaired adaptive immunity in diabetic animals

One of the challenges of studying a pathogen as virulent as SARS-CoV-2 is the requirement for ABSL-3 precautions. As such, sample neutralization by fixation or heat inactivation is required for biosafety reasons, leaving extremely limited options for downstream analysis of biological materials. We therefore adapted a method of dissociating fixed tissue into a single cell solution, originally developed for single cell RNA sequencing (Figure 3A)(17). In short, tissue was fixed in 4% PFA for 24 hours, quenched, and subsequently diced into small fragments. Tissue was then digested for 1-2 hours at 37° in Liberase TM before proceeding with standard antibody staining (Methods). We have validated that this methodology works well with flow-based immunophenotyping of common markers including CD45, CD4, CD8, B220, *et cetra* (Figure 3B), though we note that given variability in tissue digest, we have greater confidence in percentage values rather than total cell numbers using this protocol. Excitingly, we also demonstrated that this method also works well even with tissue cryoembedded in OCT (Supplementary Figure 3, Methods).

**Figure 3.**
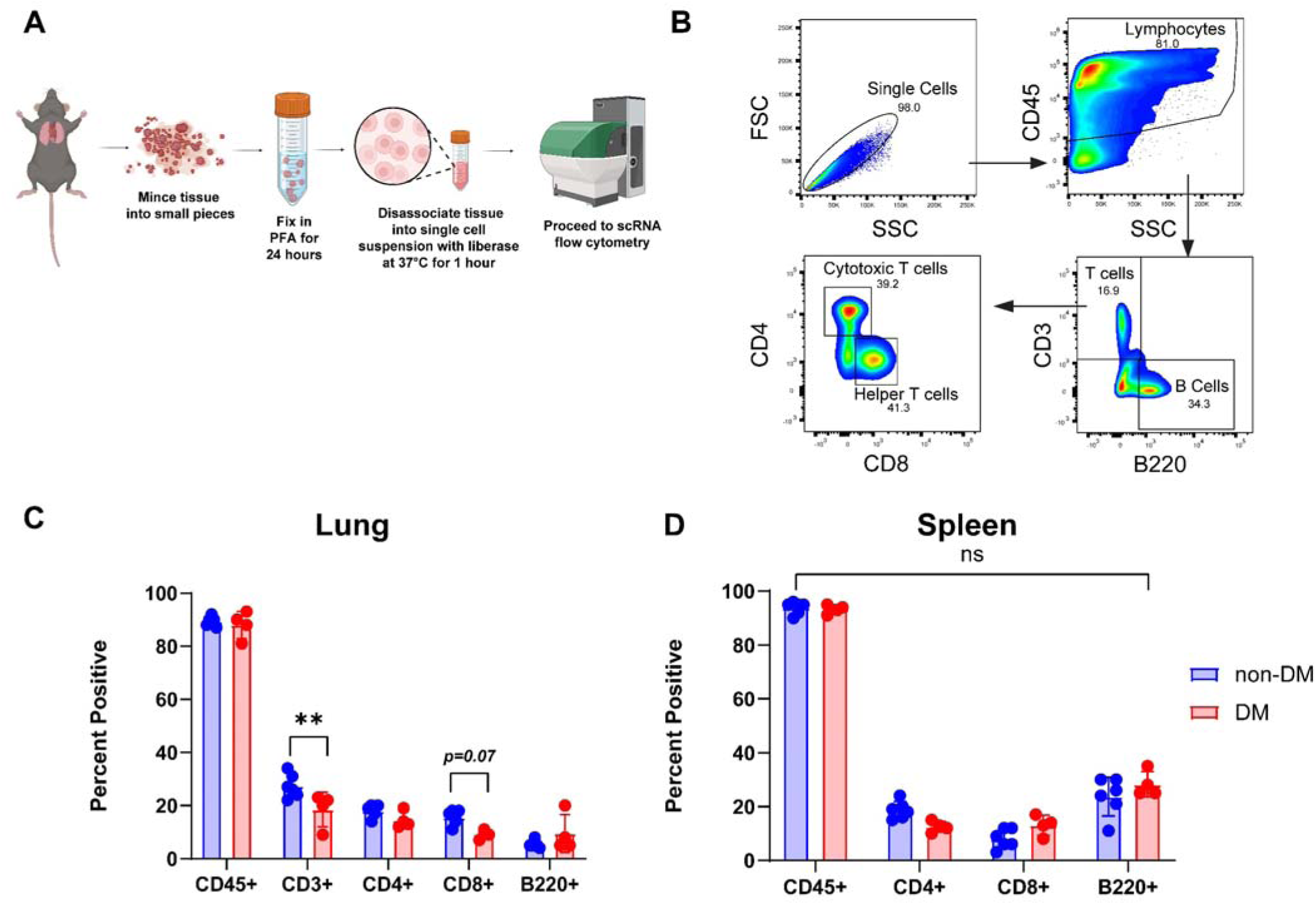
Novel technique for single cell digestion of fixed tissue and flow cytometric analysis. A. Schematic of protocol for digesting fixed tissue from ABSL-3 facility into single cell digest for flow cytometry analysis. Figure made with Biorender. B. Representative gating scheme for identification of lymphocytes (CD45^+^), all T-cells (CD3^+^), including helper T-cells (CD4^+^) and cytotoxic T-cells (CD8^+^), and B-cells (B220^+^) by flow cytometry in the lungs and spleens of non-DM and DM mice infected with MA30SARS-CoV-2. C-D. Quantification of the proportion of different immune populations in the (C) lungs and (D) spleens of non-DM and DM mice at 7 days post infection. Data in (C) and (D) are presented as Mean ± SEM, calculated using one-way analysis of variance (ANOVA), followed Sidak’s multiple comparison test. **p < 0.01.

We then went about performing flow cytometry to better characterize the differences in immune response between the T1DM and non-DM cohorts. In addition to looking at immune cell numbers from the lungs of these mice, we also compared the immunoprofile of the spleens as a representative of the peripheral immune system to assess whether treatment with STZ in any way “primed” the immune system prior to SARS-CoV-2 infection that may explain the findings from whole lung RNA seq.

Focusing especially on changes in B and T cells as suggested by our RNA-Seq findings, this approach allowed us to utilize traditional FACS-based methods to validate transcriptomic observations. This approach revealed a significant decrease in the CD3^+^ T cell population in the lungs of DM mice as compared to their euglycemic cohort and a nearly significant decrease specifically in CD8^+^ T cells (Figure 3C), consistent with the results of the whole lung bulk-RNA seq experiment. There was however no significant difference in the % of total CD45^+^, B220^+^, CD4^+^, CD8^+^ cells in the spleens of the two groups (Figure 3D), suggesting that STZ has no obvious effect on the peripheral adaptive immune system composition.

### Single-cell transcriptomic analysis reveals a distinct macrophage population in the lungs of diabetic mice post-SARS-CoV-2 infection

Recognizing that bulk RNA-Seq suffers from potential obfuscation of discrete signals driving the antiviral response from a rare cell population or cell type, we sought to better characterize the effects of hyperglycemia and T1DM on the pulmonary host response to SARS-CoV-2 by performing single-cell RNA sequencing (scRNA seq) on lungs harvested from mice infected with SARS-CoV-2. Single cell suspensions were created from the fixed lung tissue of DM and non-DM mice infected with SARS-CoV-2 at day 7 post infection as in our flow cytometry experiments. Libraries were created using the 10x Genomics Chromium Fixed RNA Profiling kit and sequenced on an Illumina NextSeq 2000. A total of 10,726 cells were sequenced with 3,197 cells from DM lungs and 7,529 cells from non-DM lungs. Cluster analysis in Seurat identified 16 separate and distinct clusters of cell types based on marker gene expression (Figure 4A). These 16 clusters include subtypes of cells found in the epithelial, mesenchymal, endothelial, and immune compartments of the lung, as was expected. Analysis of the clusters in the uniform manifold approximation and projection (UMAP) by cells labeled by sample of origin reveals a clear reduction in number of cells involved in adaptative immunity in the lungs of the DM mice (Figure 4B). In particular, the T-and B-cell clusters are overwhelmingly represented by cells from the non-DM sample. Alveolar fibroblasts seem to be more prevalent in the lungs of DM mice, possibly as a result of increased inflammation and inflammatory signals from the persistence of viral particles.

**Figure 4.**
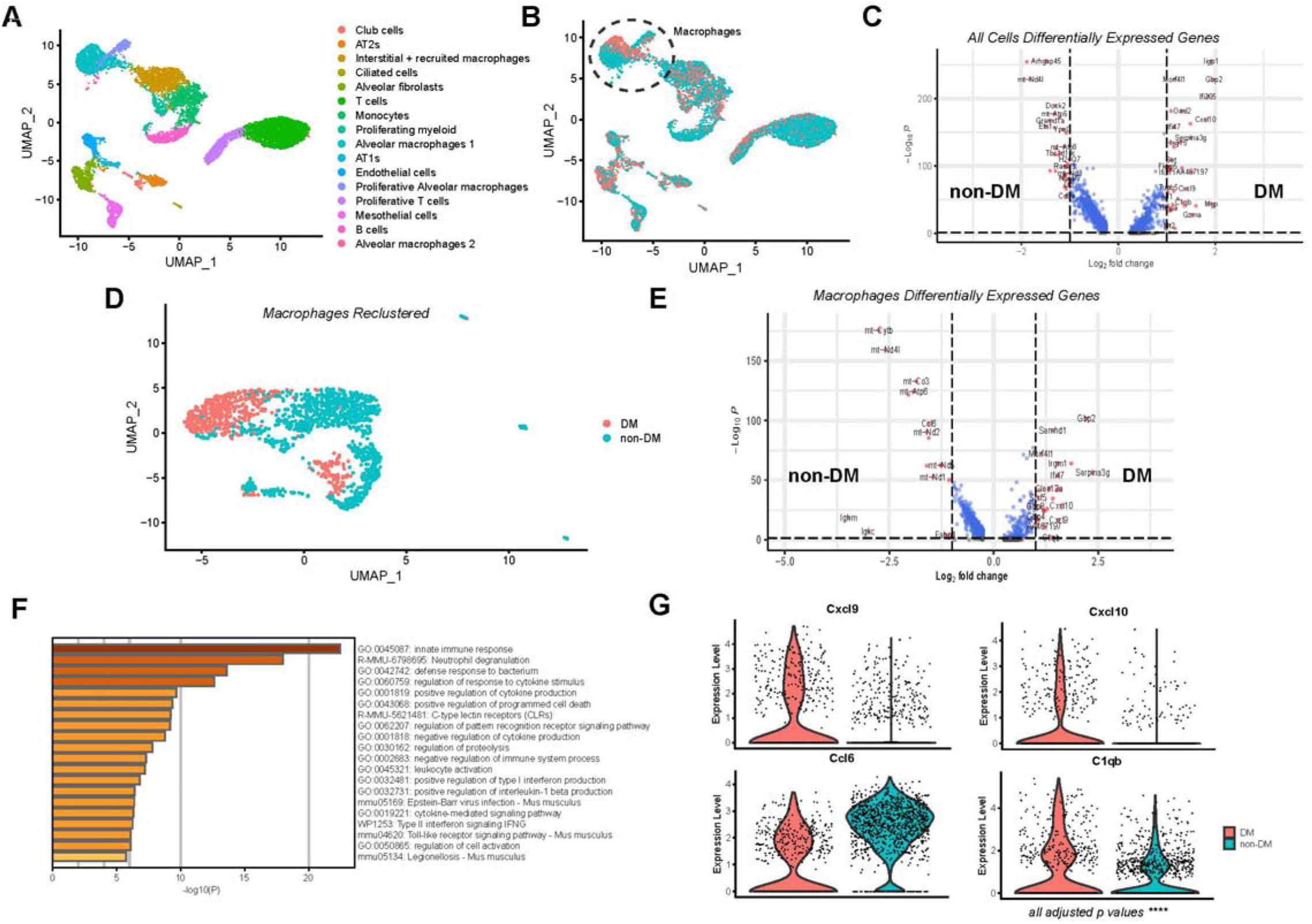
Single-cell transcriptomic analysis reveals a distinct macrophage population in the lungs of diabetic mice post-SARS-CoV2 infection. A. scRNA-seq analysis for lungs harvested from non-DM and DM mice infected with MA30-SARS-CoV-2 at day 7 post infection, cells are color coded for population clustering and presented in UMAP plot. B. UMAP plot from (A) with cells color coded by sample of origin (non-DM versus DM). C. Volcano plot of significantly differentially expressed genes in the lungs of non-DM and DM mice at day 7 post infection (differential gene expression meeting significance = 1996 genes). D. UMAP plot configured from a subset of (A) looking at macrophages with cells color coded by sample of origin (non-DM versus DM). E. Volcano plot of significantly differentially expressed genes in the alveolar macrophages of non-DM and DM mice at day 7 post infection (differential gene expression meeting significance = 1875 genes). F. The GO terms of differentially expressed genes in alveolar macrophages of non-DM and DM mice lungs infected with MA30-SARS-CoV-2 reveals DM macrophages upregulate genes responsible for inflammation. G. Violin plots of cytokine genes in alveolar macrophages compared between of non-DM and DM mice lungs infected with MA30-SARS-CoV-2. All comparisons shown in (G) have an adjusted p-value of < 5x10^-4^ using Wilcoxon Rank Sum test within Seurat.

Notably, we observed a clear difference in alveolar macrophages between the two samples (Figure 4B, circle) that did not appear to simply be a reflection of differential cell numbers. Instead, the transcriptional differences between the alveolar macrophages of the two groups were apparent in the separation in clustering on the UMAP. These differences were emphasized upon isolation and reclustering of the macrophages (Figure 4D), where the alveolar macrophages from DM mouse lungs showed almost no spatial overlap with those from non-DM mice. The transcriptional differences driving this separation are highlighted in volcano plots of significantly differentially expressed genes between these two macrophage populations (Figure 4E). Metascape analysis of the 200 most significantly upregulated genes in the DM macrophages revealed that a majority of these genes that are upregulated are involved in cytokine release and response, pro-inflammatory signals, and programmed cell death (Figure 4F, G). This heightened inflammatory state corroborates the increased weight loss and delayed recovery the DM mice experience with SARS-CoV-2 infection and may be a compensatory, yet dysregulated, response to the deficient adaptive immune response as evidenced by the lower number of adaptive immune cells seen both by flow cytometry and scRNA seq. Taken together, these findings suggest a model in which T1DM / hyperglycemia causes adaptive immune defects (especially in T cell activation / recruitment), resulting in alveolar macrophages adopting a compensatory proinflammatory phenotype and contributing to greater clinical symptoms of disease.

## DISCUSSION

Despite the start of the pandemic being December 2019 and the US government ending the federal Public Health Emergency for COVID-19 status(18–21), as of January 2024, more than one thousand Americans die from COVID per week(22)—not including those who have died from COVID-related long term health detriments. COVID-19 is thus clearly still a major health concern. Patient responses to SARS-CoV-2 infection are varied, with patient comorbidities acting as major contributors to varied outcomes.

Focusing on one such major comorbidity, we assessed whether pharmacological induction of T1DM would increase the severity of lung injury in a murine model of COVID-19 pneumonia. In spite of the multitude of studies looking at host response to SARS-CoV-2 since the pandemic began in 2019(23–25), to our knowledge no existing studies have examined T1DM as a co-morbidity specifically with SARS-CoV-2 mouse models . Our findings indicate that T1DM mice exhibit deficiency in the activation, recruitment, and function of adaptive immune cells, resulting in impaired viral clearance and a dysregulated, pro-inflammatory response of the innate immune system.

It is a widely held belief that people with DM are more vulnerable to infectious pathogens(26–28). While there have been no conclusive studies defining mechanisms behind this phenomenon—and in all likelihood, such mechanisms are multifactorial and rely equally on both the pathogen and host status—several hypotheses have been explored(28–31). Differential pathogenesis may result from the hyperglycemic environment altering antigen recognition sites (e.g. surface protein glycosylation) of pathogens(7, 8), allowing them to evade immune recognition. More recent studies have shown that glucose metabolism is critical to the function of innate immune cells in the lung and that the hyperglycemic state in patients with DM upsets this delicate balance, leaving the host more vulnerable to infections with influenza A in particular(16). Our findings recapitulate the notion that hyperglycemia and DM alter the function of the adaptive immune response with a significant deficiency in the recruitment and function of T-and B-cells, but also revealed a pro-inflammatory innate immune profile compared to the euglycemic controls. Single cell transcriptomics revealed that this proinflammatory switch was particularly apparent in alveolar macrophages. We postulate that this phenotypic change in macrophages represents a compensatory mechanism in response to impaired viral clearance by the diminished adaptive immune response, though the differential signals responsible this change are uncertain. Furthermore, the fundamental mechanism by which T1DM impairs adaptive immunity will require substantial additional research, but give the stark differences in clinical course, our work highlights the critical importance of further elucidating how T1DM and hyperglycemia so profoundly disrupt adaptive immune responses.

We must note that in contrast to recently published work(16), we did not observe a differential course of clinical disease upon H1N1 infection of T1DM animals, whereas the differences were stark and highly replicable with MA30-SARS-CoV-2 infection. Many possible reasons for this discrepancy are possible. First, there could be an outsized role of the microbiome in these responses; the gut flora is known to be highly dependent on animal housing conditions which often differs greatly between institutions. Differential dosing with virus or streptozotocin may also make an impact. Further, while DCs are not highly represented in our single cell data, we did not note obvious transcriptomic differences in these cells between T1DM and euglycemic mice upon SARS-CoV-2 infection. Future work should explore these differences.

Our study has shown that, in addition to a diminished adaptive immune response, the innate immune presence in the infected lungs of DM animals is more inflammatory compared to that in euglycemic counterparts. We posit that this is a compensatory, if ineffective, reaction to persistence of virus, but this does contradict another wildly held belief that DM is in fact protective against the development of ARDS(32–34). This prior argument posits that DM decreases the release of cytokines and interleukins, resulting in less chemotaxis and inflammation, which minimizes the risk of the dysregulated

ARDS response that leads to leaky capillaries, dysfunctional epithelium, impaired gas exchange, and eventual death(32, 34). Our findings in this model of SARS-CoV-2 infection in T1DM mice directly contradict those hypotheses and appear supported by more recent clinical data indicating that patients with T1DM and T2DM were nearly three times more likely to die in the hospital compared to their age matched peers at the beginning of the pandemic(4, 5).

In addition to the scientific advancements this study has provided, we highlight several methodological advances. Our adaptation of a technique that allows for the digestion of fixed tissue into a single cell suspension that can be used for flow cytometry should enable expanded investigation of potentially lethal pathogens, which are otherwise hindered by the requirements of high biosafety level containment. Furthermore, the ability to disaggregate fixed tissue that has been embedded (in OCT or other materials) enables the study of banked tissue specimens, opening up a range of possibilities of further inquiry with previously disregarded samples.

T1DM is a disease that largely affects children and pneumonia is the number one cause of hospitalization in children less than 5 years of age, with respiratory viruses accounting for up to 66% of those cases(9). Although we did not look into T2DM in this study, T2DM has been identified as a serious global health threat by the World Health Organization, and the worldwide prevalence of T2DM is estimated to increase to 12.2% by 2040(35). Given the new age of global viral pandemics and the increased morbidity and mortality patients with diabetes have already experienced with COVID, we can anticipate there will be future threats to the health and safety of these people in the coming decades. Further studies into the interplay between DM and immune dysfunction will uncover underlying mechanisms by which we can better adapt our treatment of an ever-growing patient population. Our results herald significant expansions in the understanding of how DM renders patients more vulnerable to an ever-present, evolving deadly threat and these findings have high potential to define new therapeutic targets to decrease viral pneumonia / ARDS severity and increase survival in these patients.

## ACKNOWLEDGEMENTS

We thank the Penn Cytomics & Cell Sorting Shared Resource Library, the Center for Host Microbe Interactions (Daniel Beiting and Dan Cutillo), and the Animal Biosafety Level 3 Core (Kellie Jurado and Peter Hewins) for their assistance in performing these studies.

## AUTHOR CONTRIBUTIONS

S.K.G. designed experiments, performed data acquisition and analysis, and manuscript preparation. G.Z., S.A.T., J.W., M.E.G., N.P.H., M.M., M.S., X.L., and A.I.W. performed data acquisition and analysis. A.E.V. oversaw experimental design and manuscript preparation.

---

**Supplemental Figure 1.**
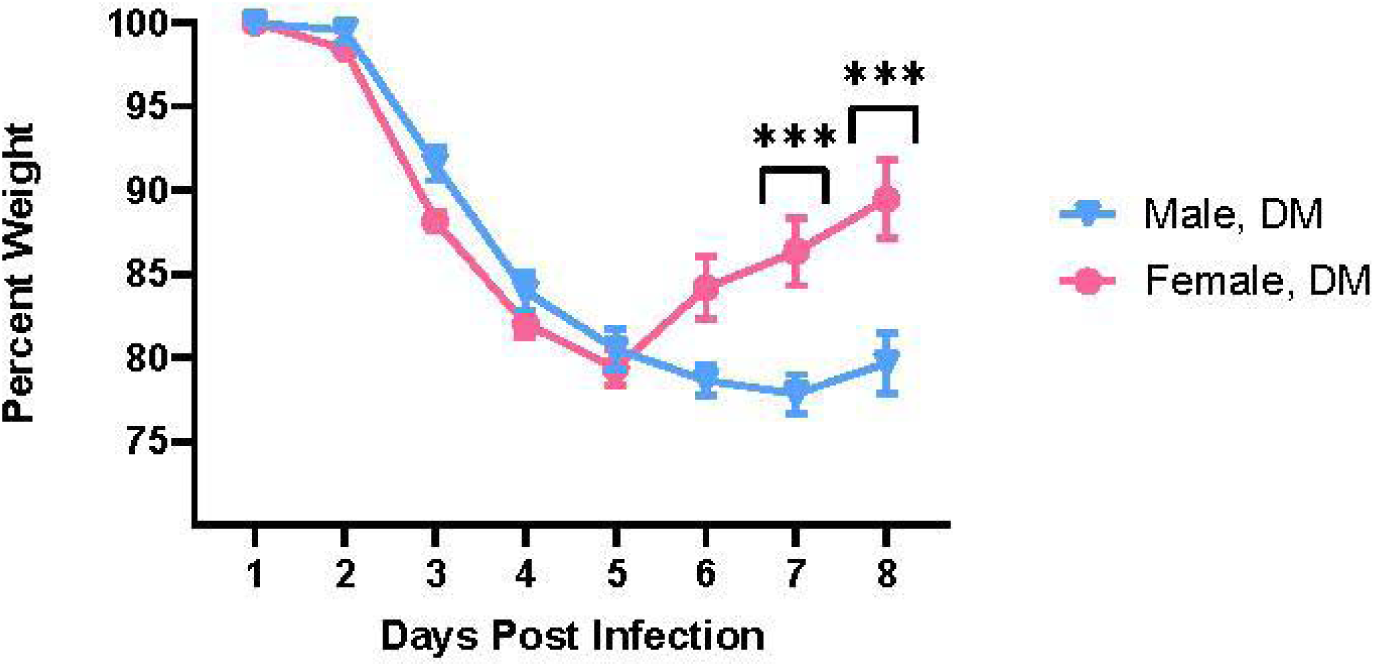
Male mice did worse in terms of weight loss and clinical illness regardless of glycemic status compared to their female equivalents. Shown here is a comparison of the time course of changes in body weight between male and female DM mice following infection with MA30-SARS-CoV-2, n = 6 to 8 per group. Data are presented as Mean ± SEM, calculated using one-way analysis of variance (ANOVA), followed Turkey’s multiple comparison test. ***p < 0.001.

**Supplemental Figure 2.**
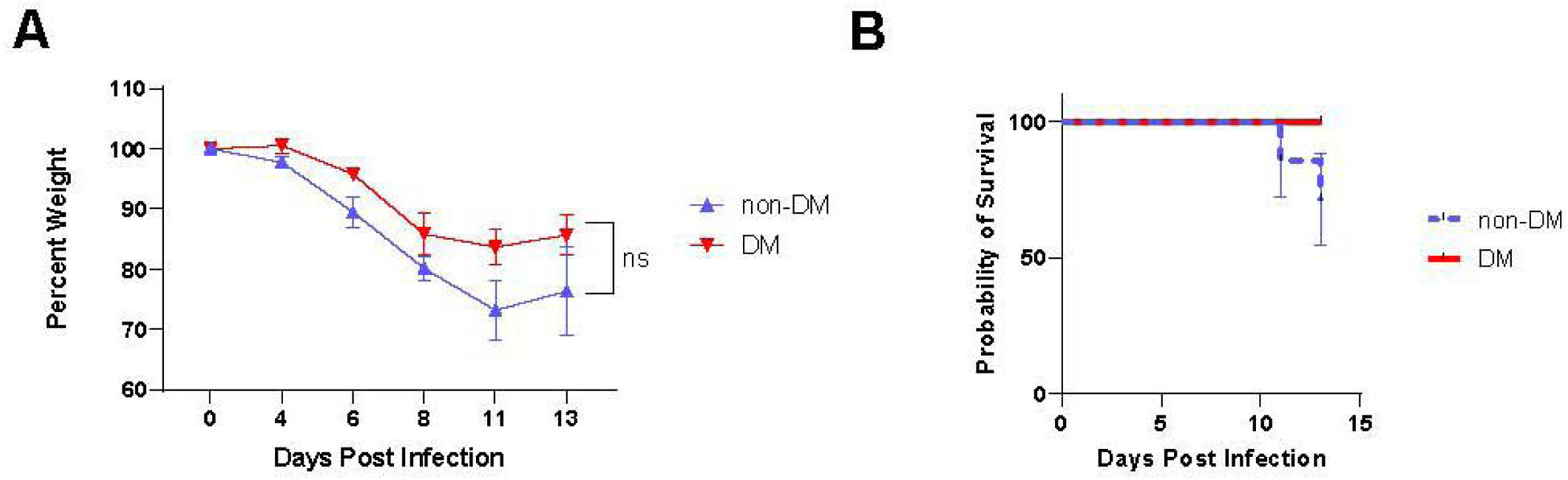
Diabetes does not confer increased susceptibility for more severe influenza infection with H1N1. A. Time course of changes in body weight in male non-DM and DM mice after H1N1 influenza (PR8) infection, n = 4 to 7 per group. B. Kaplan-Meier survival curves for non-DM and DM mice after influenza infection, not significant by log-rank test.

**Supplemental Figure 3.**
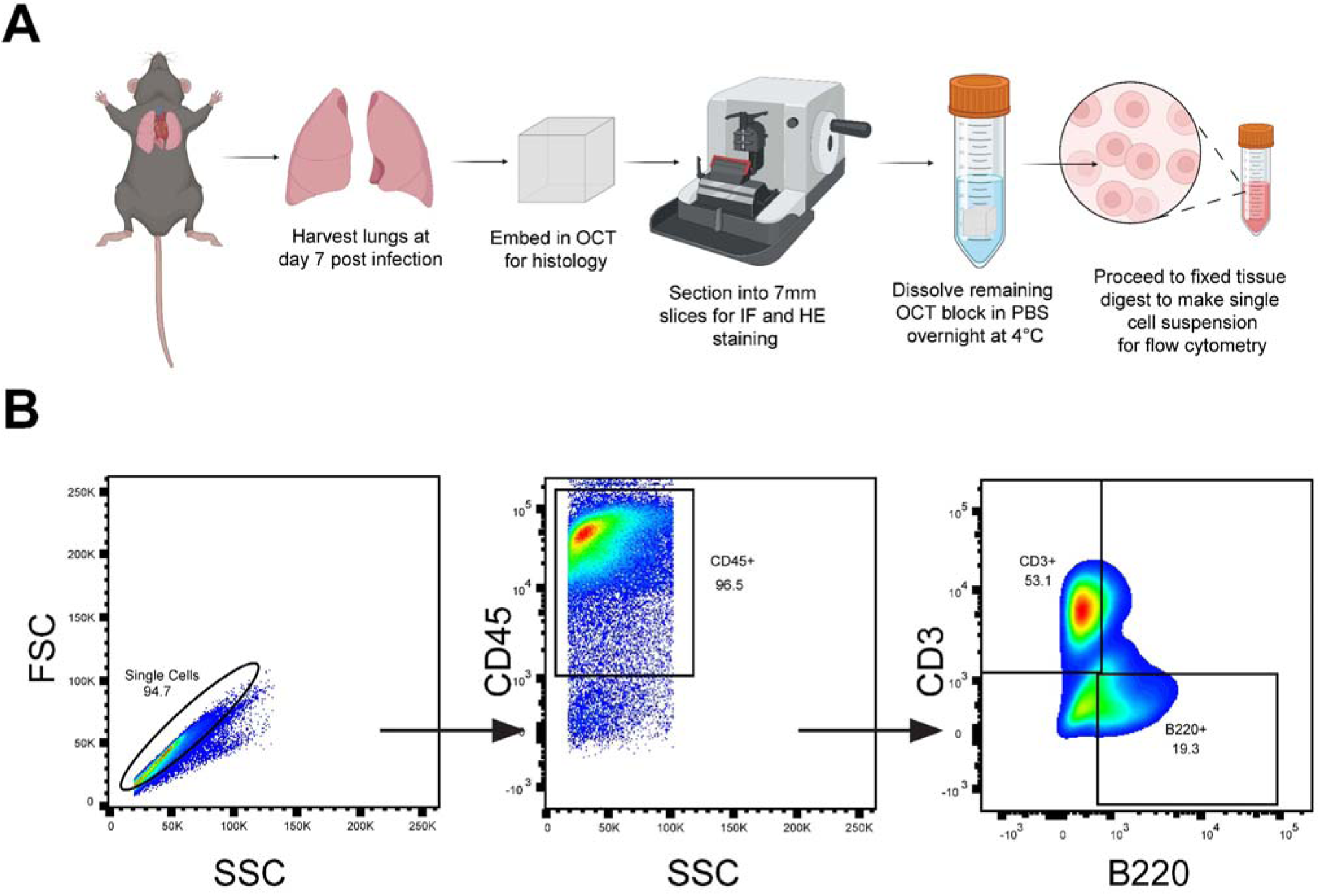
A. Schematic of protocol for digesting fixed tissue embedded in OCT into single cell digest for flow cytometry analysis. Figure made with Biorender. B. Representative gating scheme for identification of lymphocytes (CD45^+^), all T-cells (CD3^+^) and B-cells (B220^+^) by flow cytometry in the lungs and spleens of non-DM and DM mice infected with MA30SARS-CoV-2.

